# Sweet taste receptor cells may participate in mucosal immune surveillance

**DOI:** 10.1101/2022.04.28.489835

**Authors:** Yumei Qin, Xin Zheng, Shiyi Tian, Robert F. Margolskee, Sunil K. Sukumaran

**Affiliations:** School of Food Science and Bioengineering, Zhejiang Gongshang University, Hangzhou, Peoples Republic of China; State Key Laboratory of Oral Diseases & National Clinical Research Center for Oral Diseases, Department of Cariology and Endodontics, West China Hospital of Stomatology, Sichuan University, 610041, Chengdu, China; Monell Chemical Senses Center, Philadelphia, PA 19104; Department of Nutrition and Health sciences, University of Nebraska- Lincoln, Lincoln, NE 68583

## Abstract

The oral microbiome is second only to its intestinal counterpart in diversity and abundance, but its effects on taste cells remains largely unexplored. Using single cell RNASeq, we found that mouse taste receptor cells (STRCs) have a gene expression signature reminiscent of Microfold (M) cells, a central player in immune surveillance in the mucosa associated lymphoid tissue (MALT) such as those in the Peyer’s patch and tonsils. Administration of Tumor Necrosis Factor Ligand Superfamily Member 11 (TNFSF11, also known as RANKL), a growth factor required for differentiation of M cells dramatically increased M cell proliferation and marker gene expression in the taste papillae and in cultured taste organoids from wild type (WT) mice. Taste papillae and organoids from knockout mice lacking *Spib* (*Spib^KO^*), a RANKL-regulated transcription factor required for M cell development and regeneration on the other hand, failed to respond to RANKL. Taste papillae from *Spib^KO^* mice also showed reduced expression of NF-κB signaling pathway components and proinflammatory cytokines and attracted fewer immune cells. However, lipopolysaccharide-induced expression of cytokines was strongly upregulated in *Spib^KO^* mice compared to their WT counterparts. Like M cells, STRCs from WT but not *Spib^KO^* mice readily took up fluorescently labeled microbeads, a proxy for microbial transcytosis. The proportion of STRCs and other taste cell subtypes are unaltered in *Spib^KO^* mice; however, they displayed increased attraction to sweet and umami taste stimuli. We propose that STRCs are involved in immune surveillance at the taste papillae and tune their taste responses to microbial signaling and infection.

## Introduction

In mammals such as the mouse, most taste buds are found in three types of taste papillae on the dorsal surface of the tongue (1, 2). Among them, the fungiform papillae (FFP) located on the anterior tongue each house a single taste bud, and the foliate (FOP), and circumvallate (CVP) papillae located laterally and medially on the posterior tongue host a few hundred taste buds each (1, 2). The taste buds in CVP and FOP are arranged around trenches that extend down into the tongue. Each taste bud contains ~50–100 mature receptor cells classified as type I, type II, type III, and type IV cells based on morphology and marker gene expression (1–3). These cells are further classified into functional subtypes that respond to basic taste qualities of sweet, bitter, umami (subtypes of type II cells) and sour, and salt (mostly subtypes of type III cells) (1–3). Taste cells have a half-life ranging from 8-24 days and are continually regenerated from a population of stem cells located at the base of taste papillae (4–7).

16s RNA sequencing and metagenomics studies have shown that the oral cavity including the tongue dorsum is colonized by a diverse array of microbial species (8–11). Unlike cells in the non-taste lingual epithelium, taste cells are continually exposed to the oral microbiota through their microvilli that project to the surface through taste pores. However, their effects on taste signaling and taste cell regeneration have not received the deserved level of attention. The taste papillae are patrolled by a diverse population of immune cells, mostly dendritic cells including the Langerhans cell subtype, T cells and macrophages (12, 13). Most oral mucosa-resident microbes are harmless or beneficial, and the host develops immune tolerance towards them (14). However, oral dysbiosis can cause taste loss or distortion, commonly seen is patients suffering from conditions such as influenza, oral thrush (candidiasis), HIV, bacterial infection, and covid-19 (15–21). Identifying the pathways underlying the finely tuned crosstalk between taste cells, the oral microbiome and epithelial immune cells will help uncover how microbiota influence taste cells in health and disease.

The most important component of adaptive immunity at the mucosae is the mucosa-associated lymphoid tissue (MALT) (22, 23). MALT are immune inductive sites that samples the luminal microbes and generate an appropriate mucosal immune response. MALT consists of lymphoid follicles containing B and T cells, that are overlaid by an epithelial layer called follicle associated epithelium (FAE) (24). Specialized epithelial cells in the FAE called Microfold cells (M cells) transcytoses luminal microbes and pass them on to antigen presenting cells (APCs) such as dendritic cells and macrophages housed in their basal pocket (25, 26). Antigen processing and presentation by APCs stimulate the B and T lymphocytes in the underlying lymphoid follicles that mount an appropriate immune response (24, 26, 27). These effector cells then migrate to other parts of the mucosae and systemically in blood. Thus, M cells play a central role in mucosal immunity. M cells express several receptors for microbes such as glycoprotein 2 (GP2), peptidoglycan recognition protein 1 (PGLYRP1) and the poliovirus receptor (PVR) that bind to and internalize luminal microbes (28–30). They have a well-developed micro-vesicular system, but poorly developed lysosomes enabling rapid transport of their microbial cargo mostly intact across the epithelium (26). M cell differentiation is induced by tumor necrosis factor ligand superfamily member 11 (TNFSF11, also called RANKL) secreted by connective tissue cells underlying the MALT epithelium (31). RANKL binds to the receptor tumor necrosis factor receptor superfamily member 11A (TNFRSF11A), which activates the non-canonical nuclear factor kappa B (NF-κB) signaling pathway to turn on the expression of early M cell marker genes (31–34). The most prominent among them is *Spib*, a transcription factor that orchestrates the later stages of M cell differentiation (35, 36). Using single cell RNASeq (scRNASeq) of GFP-labeled mouse taste cells, we found that sweet taste receptor cells (STRCs) and a few other taste cell types express several M cell marker genes (37). The expression of a subset of these genes in taste cells was confirmed using quantitative polymerase chain reaction, *in situ* hybridization and immunohistochemistry. Consistent with this gene expression profile, RANKL administration led to marked proliferation of M cells in the taste papillae and cultured taste organoids. In contrast, taste papillae and organoids from *Spib* knockout (*Spib^KO^*) mice showed reduced expression levels of M cell marker genes and impaired RANKL-dependent M cell proliferation. *Spib^KO^* mice also showed reduced expression of NF-κB signaling pathway components and proinflammatory cytokine gene expression in their taste papillae and attracted fewer immune cells to the papillae. However, lipopolysaccharide (LPS)- induced expression of cytokines was highly upregulated in *Spib^KO^* mice compared to their wild type (WT) littermates. Using a fluorescently labelled microbead uptake assay, we show that STRCs from WT but not *Spib^KO^* mice are capable of transcytosis of luminal microparticles. *Spib* ablation did not affect the proportion of taste cell subtypes in the papillae, but *Spib^KO^* mice showed increased attraction to sweet and umami tase stimuli compared to their WT littermates. Our results indicate that STRCs possess M cell like properties and might modulate taste preference in response to microbial colonization and infection.

## Results

### STRCs express M cell marker genes

To identify genes involved in mucosal immunity expressed in taste cells, we analyzed scRNA-Seq data from *Gnat3*-EGFP-expressing (*Gnat3*-EGFP +, type II), *Tas1r3*-EGFP+ (type II) and *Gad1*-EGFP+ (type III) taste cells from CVP of respective EGFP transgenic mice. Our analysis revealed that *Tas1r3*-EGFP+ (sweet and umami receptor expressing) type II cells selectively expressed several genes critical for M cell maturation and function (Table S1) (37). The expression of a subset of these genes at the mRNA and protein levels in taste cells was confirmed using molecular and histological techniques. Endpoint PCR and quantitative real-time PCR showed robust expression of M cell marker genes *Gp2, Marcksl1, Ccl9, Anxa5, Sgne1*, and *Spib* in the CVP, while they were either expressed at much lower levels or not at all in the non-taste lingual epithelium (Figure 1 A-B). *In situ* hybridization with *Spib* sense and antisense probes showed that it is expressed in the taste cells in CVP and FOP but not in FFP (Fig 1 C-E, G-I). The antisense probe was validated in tissue sections from the Peyer’s patch (PP, the MALT in intestine) (Figure 1 F, J). These results were confirmed at the protein level using indirect immunohistochemistry with an antibody against SPIB, which stained the nuclei of subpopulations of taste cells in CVP, FOP and PP, but not FFP (Figure 1 K-N). The specificity of the SPIB antibody was confirmed by lack of staining in the PP and CVP of *Spib^KO^* mice and when the primary antibody was omitted while staining the CVP of WT mice (Figure 1 O-Q).

**Figure 1:**
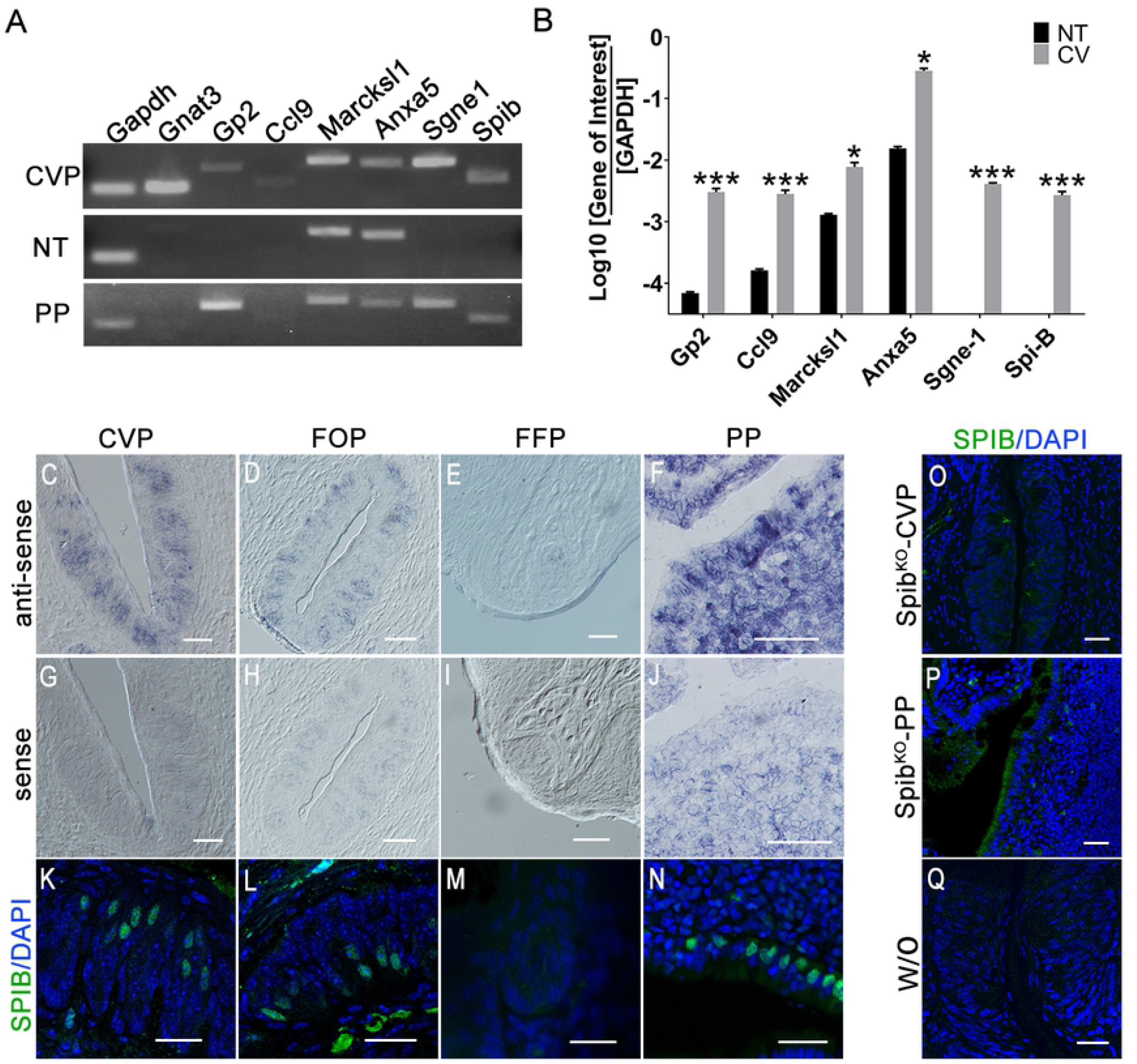
Expression of *Spib* and other M cell marker genes in taste cells. (A) End point PCR (35 cycles) of *Gapdh* (housekeeping control gene) *Gnat3* (taste tissue control gene), *Gp2, Ccl9, Marcksl1, Anxa5, Sgne-1* and *Spib* from cDNA prepared CVP, non-taste lingual epithelium (NT), and Peyer’s Patch (PP). *Sgne*-1 and *Spib* are expressed in CVP, but not in NT. *Gp2*, *Ccl9*, *Marcksl1* and *Anxa5* are expressed in both tissues, although at lower levels in the NT. (B) qPCR analysis of above genes expression in CVP and NT confirms that all M cell marker genes are highly expressed in CVP. The expression of each marker gene is plotted as the logarithm of the ratio between its cycle threshold values to those of *Gapdh*. (C-J) *In situ* hybridization using digoxigenin-labeled *Spib* anti-sense RNA probes (antisense: C-F; sense controls: G-J) produced strong signals in CVP, FOP and PP, but not in FFP. Signal from sense probe indicative of nonspecific background is much lower. (K-N) Indirect immunofluorescence confocal microscopy of cryosection from taste papillae and PP stained with an SPIB antibody similarly shows strong nuclear localization of SPIB in subpopulations of taste cells in CVP, FOP and PP but not FFP. (O-P) Absence of SPIB signal in CVP and PP of *Spib^KO^* mice proves the specificity of SPIB antibody. (Q) Omission of the primary antibody (w/o) demonstrates low nonspecific background from secondary antibody in CVP. Nuclei are counter stained with DAPI. Scale bars, 50 μm. *p<0.05, ***p<0.001.

To identify the taste cell types that express SPIB, we used double label immunohistochemistry in CVP and FOP with SPIB antibody in combination with a second antibody against a marker for all type II taste cells (TRPM5), a marker for STRCs (T1R3), a marker for bitter taste receptor expressing subset of type II cells (in the CVP and FOP, GNAT3), the type III marker CAR4 or the type I marker ENTPD2 (Figure 2). Our results confirmed that SPIB is specifically expressed in STRCs; about 91% - 97% of SPIB-expressing cells co-express TRPM5 and T1R3 in the CVP and FOP (Table S2). SPIB is weakly co-expressed (8-28%) with GNAT3, and negligibly (4-6%) with CAR4. SPIB appears to be not co-expressed with ENTPD2, although this was not quantified because type I cells wrap around other taste cells, making accurate determination of coexpression difficult.

**Figure 2.**
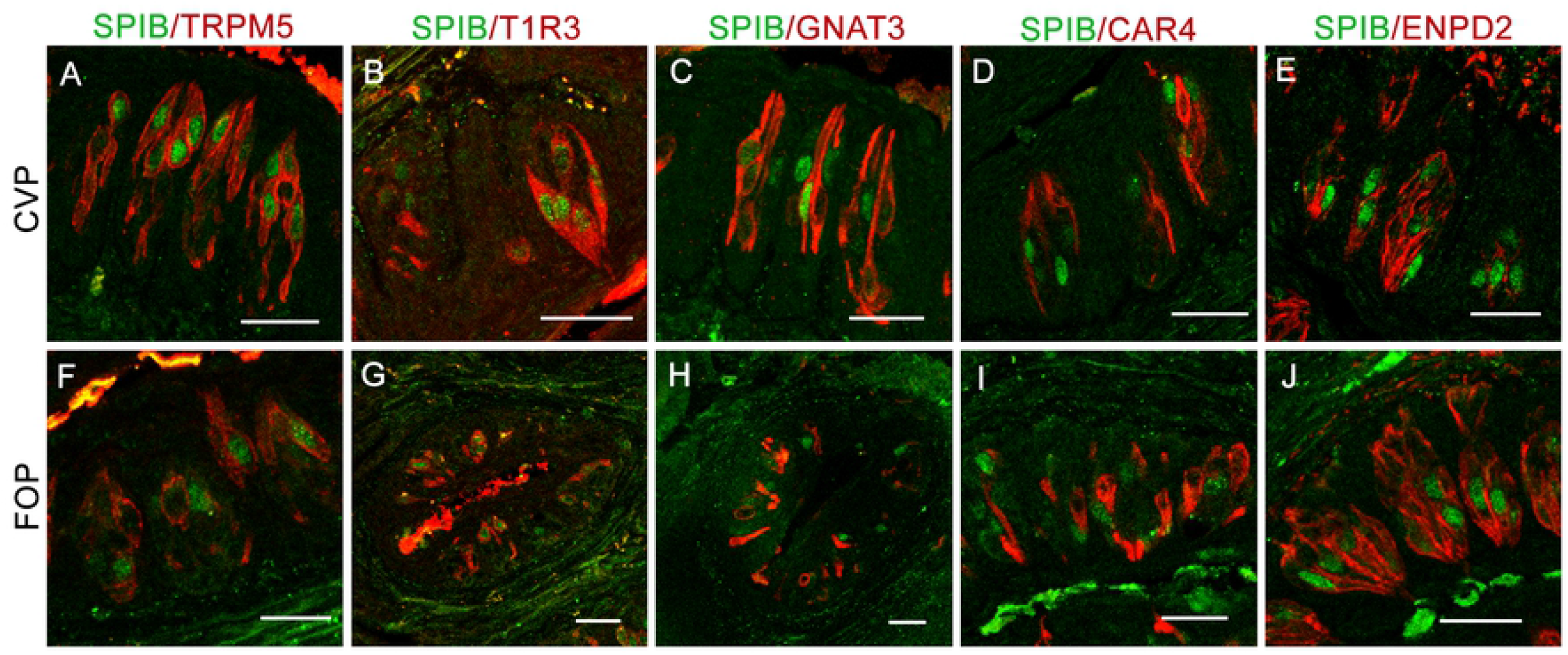
SPIB is coexpressed with T1R3. Double-labeled immunofluorescence confocal microscopy of CVP section with antibodies against SPIB and type II taste cell markers TRPM5 (A, F), T1R3 (B, G) and GNAT3 (C, H), the type III taste receptor marker CAR4 (D, I) and the type I taste cell marker ENTPD2 (E, J) show frequent co-expression of SPIB with T1R3 and less frequently with TRPM5 and GNAT3, and negligible coexpression with CAR4 and ENTPD2. Nuclei are counterstained blue with DAPI. Scale bar, 50 μm.

Double label immunohistochemistry also showed that two other M cell markers, GP2 and CCL9 appear to be coexpressed with the type II cell marker TRPM5, but not with the Type III cell marker CAR4 in the CVP, although the coexpression was not quantified (Figure 3). Of note, the strong GP2 staining at the apex of taste buds, presumably in taste cell microvilli, is similar to the pattern observed in M cells in other MALT for this receptor protein (Figure 3 C, F) (28). CVP from *Pou2f3* knockout mice that lack all type II taste cells including STRCs do not express CCL9, GP2 and SPIB (Figure S1)(38). Collectively, the results above conclusively show that STRCs express several M cell marker genes.

**Figure 3.**
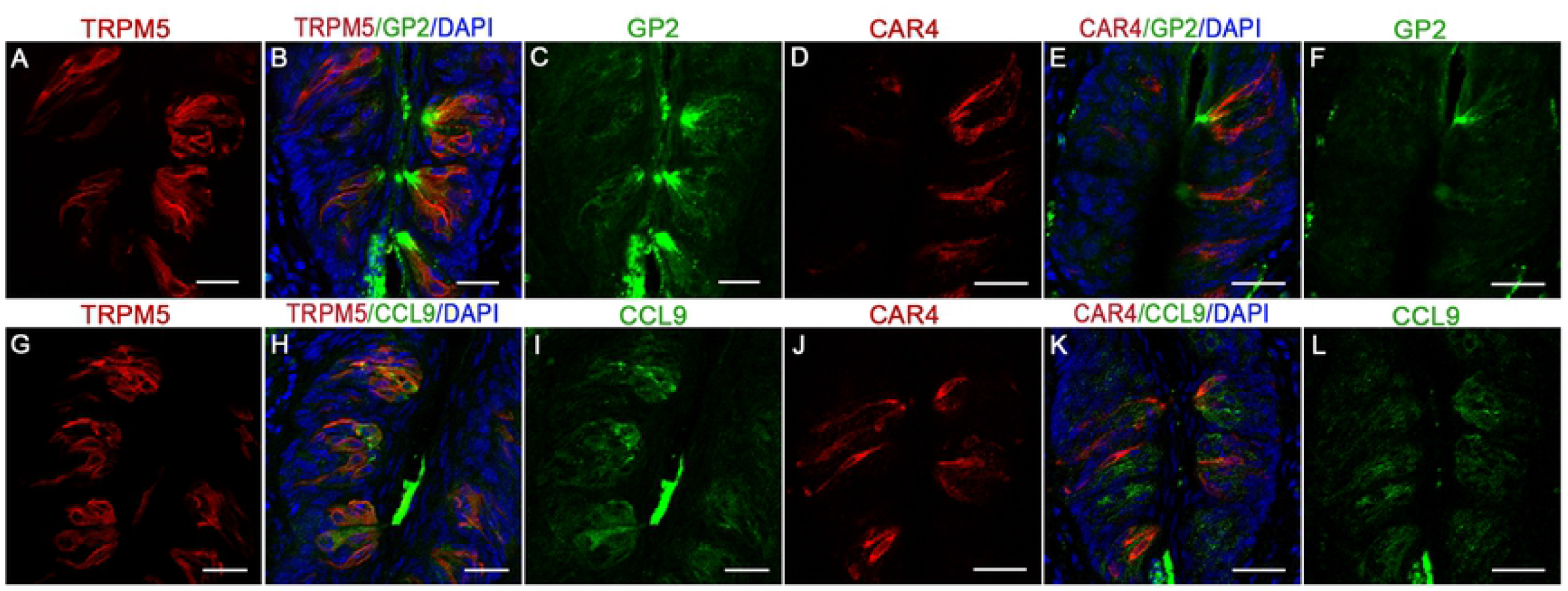
Coexpression of M Cell and type II taste cell marker proteins. Double-labeled immunofluorescence confocal microscopy of CVP sections with antibodies against M-cell markers GP2 and CCL9, with the type II taste cell marker TRPM5 (A-C and G-I) or the type III taste receptor marker CAR4 (D-F and J-L) in the CVP. Merged images show GP2 and CCL9 are coexpressed with TRPM5 (B, H), but not CAR4 (E, K). Nuclei are counterstained blue with DAPI. Scale bar, 50 μm.

### RANKL upregulates M cell marker gene expression in taste papillae and cultured taste organoids *Spib*-dependently

We next asked if RANKL can trigger *Spib*-dependent proliferation of M cells in taste papillae like it does in other MALT(31, 35, 36). The basal expression levels of M cell marker proteins GP2, CCL9, MARCKSL1 and SPIB and the mRNAs encoding *Gp2* and *Ccl9* are lower in CVP of *Spib^KO^* mice (Figure S2 A-D and I, K). Administration of RANKL led to a dramatic increase in the proportion of GP2, CCL9, MARCKSL1 and SPIB expressing cells and the levels of corresponding mRNAs in the CVP of WT but not *Spib^KO^* mice (Figure S2 E-M). Similarly, qPCR analysis showed that the expression of *Gp2, Ccl9* and *Marcksl1* mRNAs were not upregulated in the non-taste epithelium from both strains (Figure S2 M).

In the PP and intestinal villi, RANKL-treatment induces expression of various M cell marker genes in a stereotypical temporal order (35). To determine if this is conserved in taste cells, we turned to taste organoids derived from *Spib^KO^:Lgr5-EGFP* knockin or control (Lgr5-EGFP) strains. Like in PP and intestinal organoids, expression of the early M cell markers *Marcksl1* and *Ccl9* mRNAs and proteins started increasing by day 1 and reached their peaks at day 2; that of *Spib* increased gradually and reached the peak by day 3 and that of the late marker *Gp2* increased more slowly and reached its peak at day 4 after RANKL administration in organoids derived from control mice (Figure 4 A-P). Taste organoids derived from *Spib^KO^* mice, on the other hand, showed a dramatically impaired induction of *Gp2*, *Ccl9*, and *Marcksl1 mRNAs* upon RANKL administration (Figure 4 M-P).

**Figure 4.**
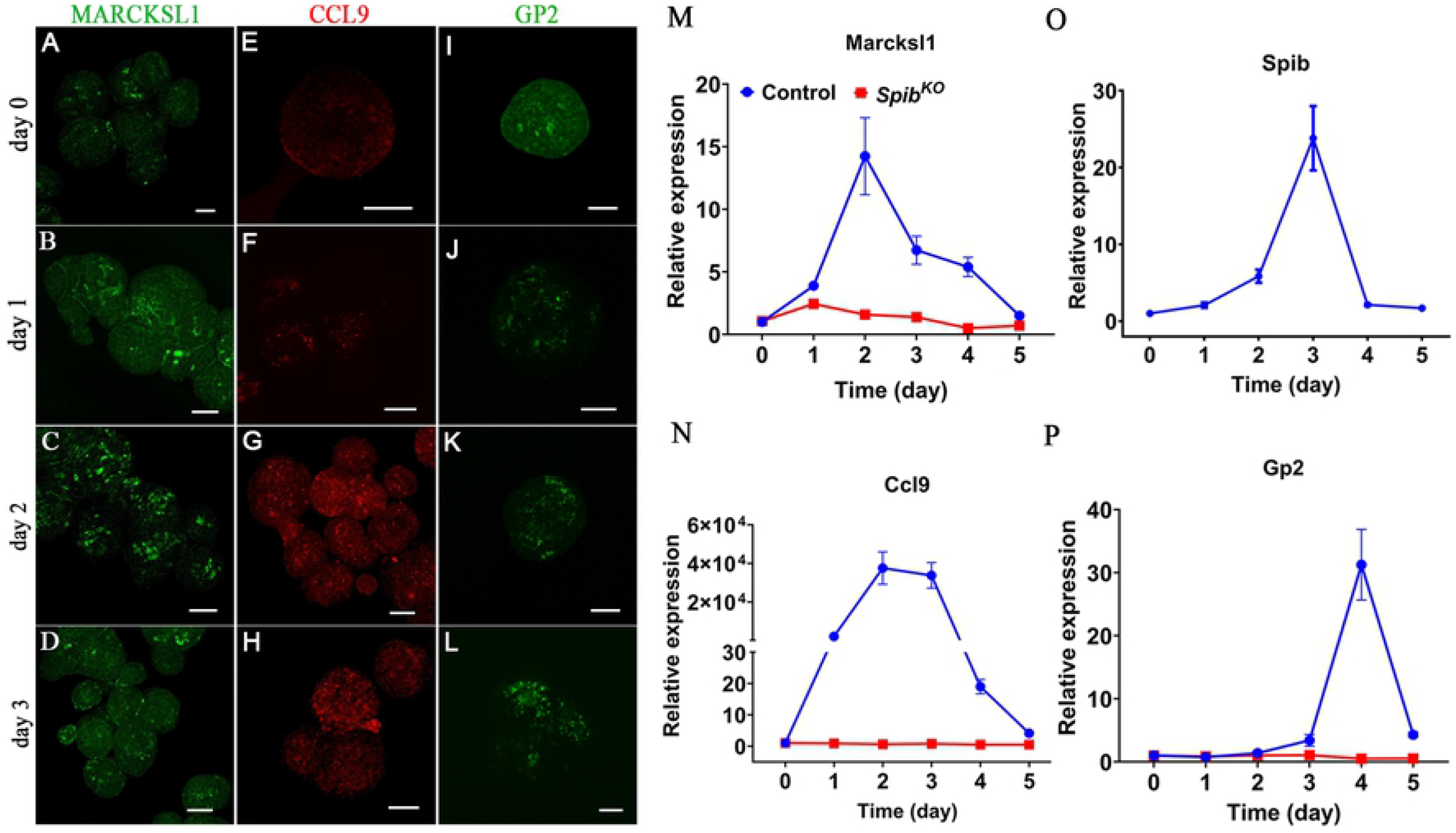
RANKL-treatment induces expression of M cell marker genes in a stereotypical temporal order in cultured taste organoids. (A-L) Indirect immunofluorescence confocal microscopy of taste organoids from control and *Spib^KO^* mice showing kinetics of expression of the M-cell markers MARCKSL1 (A-D), CCL9 (E-H) and GP2 (I-L) 0-3 days after RANKL treatment. (M-P) qPCR analysis of the expression of M-cell markers in cultured taste organoids after RANKL treatment. The kinetics of expression of GP2, CCL9 and MARCKSL1 after RANKL treatment were distinct at both protein (A-L) and mRNA (M-P) levels. *Spib^KO^* mice failed to upregulate all M cell marker genes (M-P) upon RANKL administration. Data are mean means ± SEM. Scale bars, 50 μm.

### Mucosal immune responses are impaired in *Spib^KO^* mice

Next, we compared various aspects of mucosal immunity in CVP of WT and *Spib^KO^* mice. Multiple stimuli, including RANKL, other TNF family ligands, and pathogen-associated molecular patterns such as lipopolysaccharide (LPS) signal through the NF-κB pathway (39). We asked if *Spib^KO^* mice show defects in expression of the components of the NF-κB signaling pathway and its target genes. qPCR analysis showed that the expression of the adapter protein *Myd88*, regulator proteins *Tollip, Irak3* and *Irak4* and the transcription factors *Nfkb1, Nfkb2, Rela, Rel* and *Irf6* are downregulated in *Spib^KO^* mice (Figure S3 A-B). In agreement with these findings, expression of several proinflammatory cytokines regulated by NF-κB, namely *Il1b, Il6, Mcp1* and *Tnf* is downregulated, while that of the anti-inflammatory cytokines *Il10* and *Il12* and *Ifnγ* did not change in *Spib^KO^* mice (Figure S3C). LPS is known to cause robust proinflammatory cytokine expression in CVP, and we tested if this is recapitulated in *Spib^KO^* mice (20). In contrast to baseline expression levels noted above, LPS administration triggered exaggerated cytokine gene expression in CVP of *Spib^KO^* mice compared to WT littermates (Figure S4). Next, we asked if the reduced baseline cytokine expression in *Spib^KO^* mice affected the recruitment of immune cells to the CVP. Immunostaining with antibodies against CD11B and CD45, that label a broad spectrum of immune cells or the T-Cell marker CD3 showed that *Spib^KO^* mice had fewer immune cells patrolling the CVP (Figure 5 A-F). Finally, we tested if STRCs are capable of microbial transcytosis and if this is impaired in *Spib^KO^* mice. The uptake of fluorescently labeled microbeads, a proxy for transcytosis is evident in STRCs and neighboring cells in CVP from WT, but not *Spib^KO^* mice (Figure 5 G-I).

**Figure 5.**
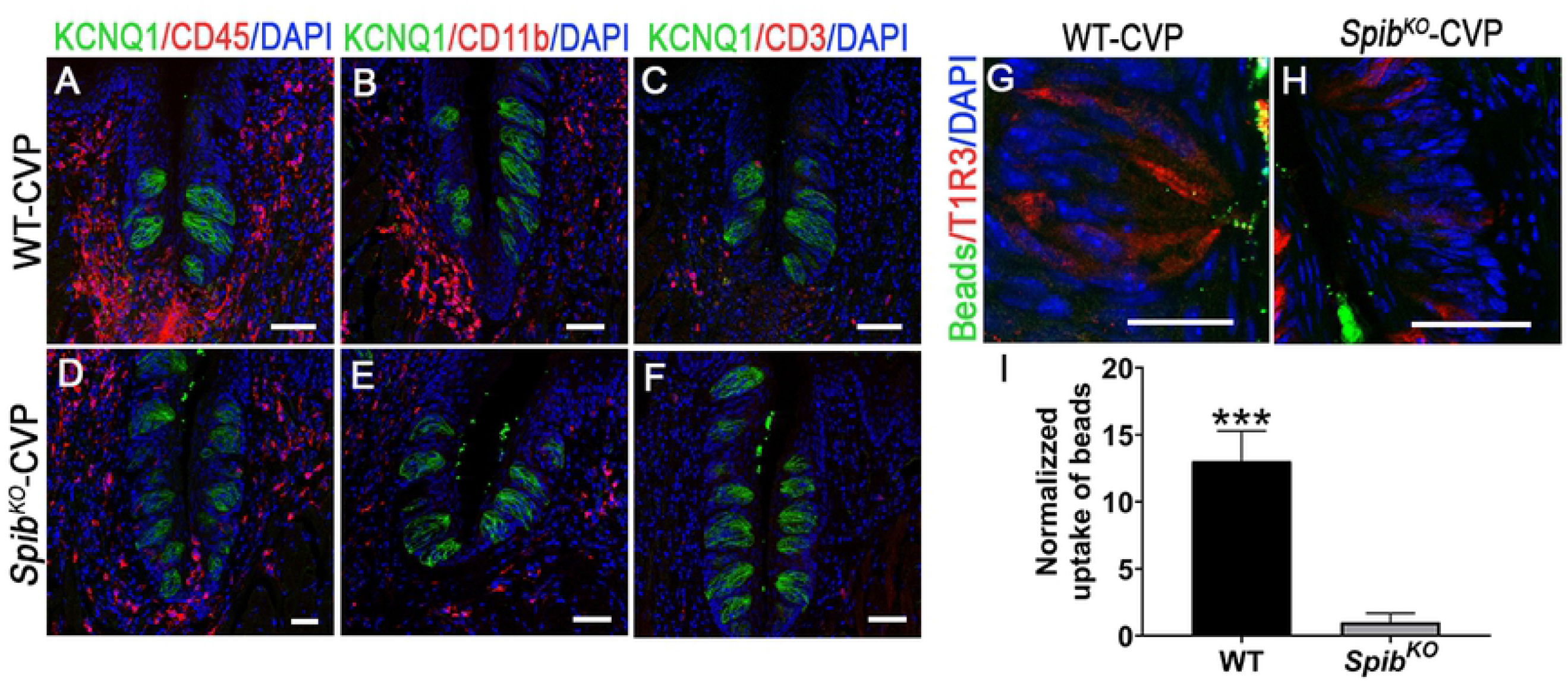
*Spib* knockout mice have impaired immune responses. (A-F) Indirect immunofluorescence confocal microscopy of CVP section from WT and *Spib^KO^* mice stained with antibodies against mouse immune cell markers CD45 (A, D), CD11B (B, E) and CD3 (C, F). Compared to WT mice, *Spib^KO^* mice had fewer immune cells in the CVP. (G-H) Uptake of 200-nm diameter fluorescent beads in T1R3 positive cells from CVP of WT (G) and *Spib^KO^* mice (H) were observed using confocal microscopy. (I) Uptake of beads was quantitated by image analysis and normalized so that the average uptake in WT mice was 1.0. Compared to WT mice, *Spib^KO^* mice uptake fewer beads. ***p<0.001. Scale bars, 50 μm.

### *Spib^KO^* mice have enhanced attraction to sweet and umami tastants

Given its role in regeneration and function of M cells, we asked if ablation of *Spib* cause defects in taste cell regeneration and/or taste preference. Immunostaining with antibodies against the taste marker proteins T1R3, TRPM5, GNAT3 and CAR4 showed that the proportion of type II and III cells are unaltered in the CVP of *Spib^KO^* mice compared to WT (Figure S5A). These results were confirmed by qPCR analysis of the corresponding mRNAs (Figure S5 B-F). However, in brief access taste tests, *Spib^KO^* mice displayed increased attraction to the prototypical sweet and umami taste stimuli sucrose and monopotassium glutamate compared to WT littermates (Figure 6A, C). The attraction to sucralose appeared to be higher as well in *Spib^KO^* mice, although not statistically significant (Figure 6B). On the other hand, the responses to denatonium benzoate (bitter), sodium chloride (salty), and citric acid (sour) that are mediated by taste cell types other than STRCs were unchanged between the two strains (Figure 6 D-F).

**Figure 6.**
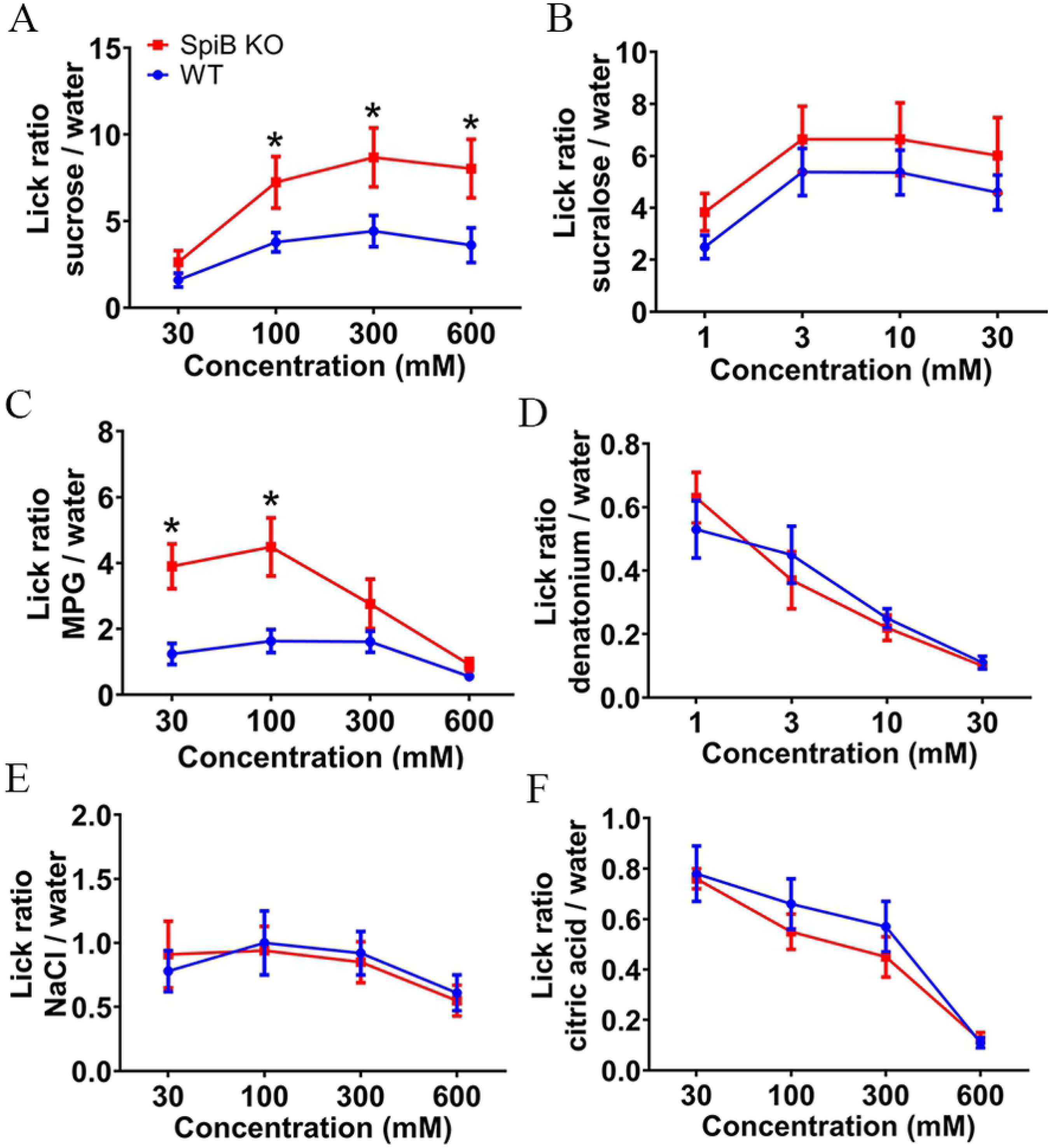
*Spib^KO^* mice show increased behavioral attraction to sweet and umami tastants. Brief access tests were used to measure behavioral responses to sweet (sucrose and sucralose, A and B), Umami (potassium glutamate [MPG], C), bitter (denatonium, D), salty (NaCl, E) and sour (citric acid, F) taste stimuli. Compared to littermate control WT mice, *Spib^KO^* mice show increased lick responses to sweet and umami (sucrose and MPG), while the response to other taste stimuli are unchanged. Lick ratios were calculated by dividing the number of licks to a taste solution by the number of licks to water in each test session. Data are means ± SEM analyzed with two-way ANOVA with post hoc t test. N= 12 (WT mice) and 14 (*Spib^KO^* mice). *p<0.05.

## Discussion

Mucosae are important routes of microbial colonization, and animals have evolved a strong mucosal immune system comprised of both innate and adaptive components to counter infection (40–42). The MALT is a type of secondary lymphoid tissue critical for mucosal adaptive immunity. The MALT in the gut (PP) and tonsils are the best studied, but MALT occurs in other mucosae such as the salivary glands, nasopharynx, conjunctiva and tear ducts as well (24, 43, 44). The mucosa in the tongue dorsum is heavily colonized by the oral microbiome, even more so than the buccal mucosa (9, 45). Most of the tongue surface is made up of keratinized stratified squamous epithelium that acts as an effective barrier to microbial invasion. However, the microvilli of taste cells project to the tongue surface through the taste pores in the taste bud and presumably represent an easier route for microbial invasion. The trenches in the CVP and FOP surrounded by taste buds are ideal sites for long-term microbial colonization, as they are largely shielded from salivary flushing. However, the effects of the oral microbiota on taste cells have not been studied in sufficient detail so far. The expression of Toll-Like receptors (TLRs), interferon receptors and their downstream signaling pathway components in taste cells has been documented (46–48). Administration of LPS or double-stranded RNA that binds to TLRs and mimic bacterial and viral infection respectively, activates TLR and interferon signaling pathways in taste cells, and diminishes taste cell regeneration and taste transduction, likely by promoting secretion of proinflammatory cytokines (20, 47, 49, 50). Similarly, knocking out *Tlr4*, the primary receptor for LPS in mice leads to diminished taste response to sugars, lipids, and umami (48). Interestingly, sweet, and bitter taste receptors bind microbial metabolites and mediate immune responses in several extra oral tissues, but whether they do so in taste papillae itself is not known (51, 52).

There are several functional and developmental similarities between cells in the taste and intestinal epithelia. For e.g., intestinal epithelial cells and taste cells in the CVP and FOP are of endodermal origin and arise from *Lgr5*-expressing stem cells (4, 53, 54). Taste receptors and other members of the taste transduction machinery are also expressed in the nutrient sensing enteroendocrine cells and the microbe- and parasite-sensing tuft cells in the intestine (55–60). Finally, both taste and intestinal epithelial cells are heavily exposed to the respective microbiota, and it is plausible that some taste cell types may share functional features of M cells. Indeed, we identified an M cell like gene expression signature in STRCs and a few other taste cell types using scRNASeq (Table S1, Figures 1–3). Strikingly, RANKL administration led to M cells proliferation in CVP and upregulation of M cell marker genes in cultured taste organoids in the same temporal order observed in the PP and PP-derived organoids (Figures 4, S2). Finally, as in the PP, M cell marker gene expression in CVP requires *Spib. Spib^KO^* mice have low basal levels of M cell gene expression and fail to respond to RANKL (Figures 4 &S2). In agreement with these observations, STRCs from WT but not *Spib^KO^* mice are able to transcytose luminal microbeads (Figure 5 G-I). Thus, all evidence indicates that STRCs have M cell like properties. However, it is not clear if STRCs are the only cells with M cell properties. For e.g., it is not clear if all the cells expressing M cell marker proteins upon RANKL stimulation are STRCs or if only STRCs are capable of transcytosis. Some type II bitter and type III taste cells also express TNFRSF11A (RANKL receptor), and it possible that they form a sizeable proportion of cells expressing M cell marker genes upon RANKL stimulation (Table S1). Similarly, although the fluorescent microbead uptake assay shows colocalization of beads in STRCs, several beads are present in unlabeled cells (Figure 5 G-I). These beads might represent a subset that were transcytosed out of STRCs or directly taken up by other taste cell types. However, it is unlikely that type I or IV cells are capable of transcytosis since their microvilli do not project out of the taste pore (3). In summary, our data indicates that STRCs likely form the bulk of M cell like taste cells, although it is possible that a smaller subset of other taste cells, likely type II bitter taste receptor cells and/or type III cells can also acquire M cell properties.

Despite the similarities outlined above, there are key differences between STRCs and classical M cells; STRCs do not have the basal pocket found in PP M cells that houses APCs. In this respect, they are analogous to villous M cells that spontaneously transdifferentiate from enterocytes in apices of intestinal villi (61, 62). Similarly, villous M cells and the taste papillae are not associated with underlying germinal centers (63). Villous M cell stimulated T and B cell maturation may occur in the intestinal lamina propria, but it is not known if this occurs in the lamina propria of taste papillae (27). However, taste papillae from WT mice contain a diverse population of immune cells, and *Spib^KO^* mice have far fewer of them, strongly supporting a role for STRCs in immune surveillance (Figure 5 A-F). Of note, although not present in rodents, humans and most other mammals have lingual tonsils located near the CVP and FOP. They contain MALT, where STRC-stimulated T and B cells may mature, although this might happen in the cervical or other lymph nodes as well. In fact, the mesenteric lymph nodes appear to be the primary sites of T and B cell maturation in intestinal mucosa (64).

What other roles might the M cell like properties of STRCs serve? Taste cells secrete cytokines such as IL-10 and TNF, which affect taste signaling (49, 50, 65). M cells are known to regulate secretion of cytokines in the PP(66). *Spib^KO^* mice express lower levels of pro-inflammatory cytokines and have fewer immune cells in the taste papillae while the expression of anti-inflammatory cytokines was unaltered (Figure 5 and S3C). The expression of components of the NF-κB signaling pathway that regulate acute phase cytokine gene expression is lower in CVP of *Spib^KO^* mice, which might underlie this phenotype (Figure S3 A-B). Of note, increased T-cell recruitment to taste papillae was observed in IL-10 knockout mice, which also had smaller taste buds and fewer taste cells per bud(65). The lower levels of immune cell recruitment in CVP of *Spib^KO^* mice likely reflects downregulation of cytokines, especially chemokines (Figures 5 A-F and S3 C). However, upon LPS stimulation, *Spib^KO^* mice show exaggerated inflammatory cytokine gene expression. These data and the inability of STRCs in *Spib^KO^* mice to transcytose luminal particles indicate that mucosal immune responses are impaired in the CVP in the absence of antigen surveillance. *Spib* is required for the development and regeneration of M cells in the major GALT tissue and plasmacytoid dendritic cells (35, 36, 67). However, the proportion of STRCs and other taste cells in *Spib^KO^* mice is unaltered, suggesting that its role in taste cells is likely restricted to regulating mucosal immunity (Figure S5). A direct role in regulating taste signaling itself remains an open question. The higher behavioral attraction to sweet and umami stimuli in *Spib^KO^* mice could arise from reduced expression of proinflammatory cytokines or changes in expression of downstream components of the taste signaling pathway (Figure 6). Our findings indicate that STRC-mediated immune surveillance is a key aspect of oral mucosal immunity that can affect taste preference and nutrition. Dysregulation of this pathway may lead to microbial infection and taste loss.

## Materials and methods

### Animals

All animal experiments were performed in accordance with the National Institutes of Health guidelines for the care and use of animals in research and reviewed and approved by the Institutional Animal Care and Use Committee at Monell Chemical Senses Center (protocols: 1163 and 1151 to RFM). Animals were housed with a 12-h light/dark cycle and open access to food and water. *Spib^KO^* mice on a C57BL/6J background were obtained from Dr. Lee Ann Garrett-Sinha lab (State University of New York, Buffalo, U.S.A) (68). Lgr5-EGFP-IRES-CreERT2 mice (stock #008875) were purchased from the Jackson Laboratory. *Tas1r3*-GFP, *Gad1*-GFP and *Gnat3*-GFP transgenic mice were as previously described(69–71). Lgr5-EGFP-IRES-CreERT2 mice were crossed with *Spib^KO^* mice to obtain Lgr5-EGFP-IRES-CreERT2:*Spib^KO^* and Lgr5-EGFP-IRES-CreERT2:*Spib^+/+^* animals for generating taste organoids. Genotypes were confirmed using primer sets recommended by Jackson Laboratory and Sinha lab.

#### RANKL preparation and injection

The expression and purification of recombinant mouse RANKL were performed as previously described with some modifications (31). Truncated mouse RANKL transcript (encoding amino acids 137-316 of mouse RANKL) was PCR amplified (forward primer: CACCCCCGGGCAGCGCTTCTCAGGAGCT, reversed primer: GAGACTCGAGTCAGTCTATGTCCTGAAC) and then cloned into pGEX-5X-2 vector (GE Healthcare) between SmaI and XhoI sites. The insert was verified by sequencing and then transformed into BL21 *Escherichia coli* strain (Stratagene) for expression. Bacteria harboring expression plasmids were selected and grown in Luria-Bertani (LB) media supplemented with 100 μg/ml ampicilin. The cultures were induced with 20 μM isopropyl -D-1-thiogalactopyranoside (IPTG) for 16 h at 25°C, and the glutathione-S transferase (GST) tagged RANKL (GST-RANKL) were purified from bacterial lysate by affinity chromatography on a GSTrap FF column (Cat. No. 71-5016-96 AM, GE Healthcare, Chicago, IL) followed by dialysis against multiple changes of phosphate buffered saline (PBS), pH 7.4. Recombinant GST used as a control was prepared by the same method using empty pGEX-5X-2. GST or GST-RANKL, with at least 95% purity as demonstrated by SDS–PAGE was administered to *Spib^KO^* and their littermate control animals by intraperitoneal injections (250 μg/day/mice for three days). Mice were sacrificed at day 4 and lingual epithelium or lingual tissue were collected.

### Isolation of lingual epithelium

Lingual epithelium was enzymatically peeled as described (37). Mice were sacrificed by CO_2_ asphyxiation, and the tongues were excised. An enzyme mixture (0.5 ml) consisting of dispase II (2 mg/ml; Roche, Mannheim, Germany; cat. no. 04942078001) and collagenase A (1 mg/ml; Roche cat. no. 10103578001) in Ca^2+^-free Tyrode’s solution (145 mM NaCl, 5 mM KCl, 10 mM HEPES, 5 mM NaHCO_3_, 10 mM pyruvate, 10 mM glucose) was injected under the lingual epithelium and incubated for 15 min at 37°C. Lingual epithelia were peeled gently from the underlying muscle tissue and used for single-cell RNA-Seq, FACS sorting, or RNA isolation.

### Fluorescence-activated cell sorting

GFP-fluorescent *Lgr5+* taste cells from Lgr5-EGFP-IRES-CreERT2; *Spib^KO^* and Lgr5-EGFP-IRES-CreERT2;*Spib^+/+^* mice were isolated by Fluorescence-activated cell sorting (FACS) as described (72, 73). Briefly, the region of the lingual epithelium containing the CVP from three mice was excised and pooled, minced into small pieces, incubated with trypsin (0.25% in PBS) for 10-25 min at 37°C, and mechanically dissociated into single cells using heat-pulled Pasteur pipettes. Cell suspensions were filtered using 70-μm cell strainers (BD Biosciences, Bedford, MA; cat. no. 352350) and then with 35-μm cell strainers (BD Biosciences cat. no. 352235). Cells were sorted into culture medium for organoid culture, based on the enhanced green fluorescent protein (EGFP) signal (excitation, 488 nm; emission, 530 nm).

### 3D taste organoid culture

Taste organoids were cultured as described (72, 73). Briefly, FACS sorted GFP fluorescent cells were mixed with 4% chilled Matrigel (v/v; BD Biosciences, San Jose, CA; cat. no. 354234) and maintained in DMEM/F12 (Thermo Fisher, Waltham, MA; cat. no. 11320-033) supplemented with HEPES (10mM, Thermo Fisher cat. No. 15630080), GlutaMAX (2 mM, Thermo Fisher; cat. No. 35050061), Wnt3a-conditioned medium (50%, v/v), R-spondin-conditioned medium (20%, v/v), Noggin-conditioned medium (10%, v/v), N2 (1%, v/v; Thermo Fisher; cat. No. 17502-048), B27 (2%, v/v; Thermo Fisher; cat. No. 12587-010), Y27632 (10 μM; Sigma-Aldrich; cat. No. Y0503), and epidermal growth factor (50 ng/mL; Thermo Fisher). Wnt3a- and R-spondin-conditioned medium were generated from Wnt3a and R-spondin stable cell lines as described (74). Noggin conditioned medium was made in house. The culture medium was changed first at day 5-7 and once every 2-3 days thereafter. After 14 days culturing, recombinant mouse RANKL was added into the fresh culture medium at concentration of 50 ng/ml for 5 consecutive days. At various time points (day 0 to day 5), taste organoids were harvested.

### PCR and qPCR

Total RNA was isolated taste papillae or NT epithelium dissected from freshly peeled lingual epithelium or taste organoids using the PureLink mini kit (Thermo Fisher; cat. no. 12183018A) with on-column DNA digestion (using PureLink DNase, Thermo Fisher; cat. no. 12185010) and converted into cDNA using Super Script VILO kit (Thermo Fisher; cat. no. 11755050). PCR and qPCR were done as previously described. All qPCR results were normalized using the ΔΔCt method with *Gapdh* as reference. Primers used are shown in Table S3.

#### Tissue Preparation

Mice were sacrificed by cervical dislocation and taste papillae-containing portions of the tongue were quickly removed and briefly rinsed in ice-cold PBS. For *in situ* hybridization, tissues were freshly frozen in Tissue-Tek O.C.T. mounting media (Sakura) using a 100% ethanol dry ice bath and then sectioned within 1 h after dissection. For immunohistochemistry, tissues were fixed for 1 h overnight at 4°C in 4% paraformaldehyde/1× PBS and cryoprotected in 20% sucrose/1 × PBS overnight at 4 °C before embedding in O.C.T. Sections (8–10 μm thickness) were prepared using a CM3050S cryostat (Leica Microsystems) and applied on precoated microscope slides (Superfrost plus; Fisher Scientific). Sections were dried at 40 °C for 20 min and immediately used for *in situ* hybridization or stored at −80 °C for immunohistochemistry.

### *In Situ* Hybridization

*In situ* hybridization of taste papillae was done as previously described (73, 75). Fresh sections were fixed for 10 min in 4% paraformaldehyde/1×PBS, permeabilized by a 10-min incubation at 37°C in 1 M Tris-HCl (pH 8.0)/0.5 M EDTA containing 10 μg/mL proteinase K (Boehringer Mannheim), postfixed for 10 min in 4% paraformaldehyde/1× PBS, and then acetylated for 10 min. All steps were followed by three 5-min washes with diethyl pyrocarbonate- (DEPC)-treated 1× PBS. Slides were then prehybridized for 1 h at room temperature in a mixture containing 50% deionized formamide, 5× saline/sodium citrate (SSC), 5× Denhardt’s solution, 50 μL/mL salmon sperm DNA, 2.5 μL/mL of yeast tRNA, and 2.5 M EDTA in DEPC-treated water. For hybridization, aliquots of the mixture were heated at 85 °C for 10 min to denature yeast tRNA, and full-length DIG-labeled *Spib* RNA probe (primer sequence, forward: TAATACGACTCACTATAGGCCTGCTCTGAACCACCAT; reverse: AATTAACCCTCACTAAAGGCGGATTTCGCCTGTCTTGGC) was added to yield a concentration of 0.3 μg/mL. RNA probe mixtures were heated at 85 °C for 3 min to denature the probe and then immediately chilled on ice. Hybrislip plastic coverslips (Invitrogen) were used to keep sections from drying out during hybridization, and slides were placed in a humidified chamber, sealed in a large moist zip-lock bag, and incubated at 65 °C overnight. Plastic coverslips were removed by soaking in 5× SSC prewarmed to 65 °C. Slides were washed three times for 30 min each time in 0.2× SSC and once for 10 min in PBS with 0.1× Triton X-100 (PBST). The slides were then blocked for 1 h at room temperature with 10% heat-inactivated normal goat serum, followed by a 3 h incubation at room temperature with anti–DIG alkaline phosphatase (1:1,000; Boehringer) in blocking solution. Alkaline phosphate labeling was detected by incubation overnight at room temperature in the dark with a nitroblue tetrazolium plus 5-bromo-4-chloro-3 indolyl-phosphate mixture (Roche) with levamisole (Sigma). Slides were washed in PBST, rinsed in water, dehydrated with an increasing series of ethyl alcohol, cleared with Histoclear (National Diagnostics), and coverslipped with Permount (Fisher Scientific). Antisense and sense RNA probes were used at equivalent concentrations and run in parallel in the same experiment to ensure equivalent conditions. For each experiment, a positive control hybridization using *Tas1r3* probe was done. The specificity and sensitivity of the *Spib* RNA probe was validated by *in situ* hybridization with Peyer’s patch tissue isolated from WT mice as positive control.

### Immunostaining

Immunostaining of sections containing CVP, FOP or FFP papillae was done as described (37, 75). Briefly, sections were rinsed in PBS and blocked with SuperBlock blocking buffer (Thermo Fisher; cat. no. 37515) supplemented with 0.3% (v/v) Triton X-100 and 2% donkey serum for 30 min at room temperature. Sections were incubated overnight with primary antibodies at 4°C. After 3×5 min washing with 1× PBS, species-specific secondary antibodies were used to visualize specific taste cell markers. 6-Diamidino-2-phenylindole (DAPI; 1:1000) in deionized water was used to visualize the nuclei following secondary antibody staining. The primary and secondary antibodies used in this study are listed in Table S4.

Immunostaining of taste organoids was done as previously described(72, 73). Briefly, cultured organoids were collected through centrifuging and fixed for 15 min in fresh 4% paraformaldehyde in 1× PBS supplemented with MgCl2 (5 mM), EGTA (10 mM), and sucrose (4%, wt/v). After 3×5 min washing with 1× PBS, organoids were blocked for 45 min with SuperBlock blocking buffer (Thermo Fisher cat. no. 37515) supplemented with 0.3% (v/v) Triton X-100 and 2% (v/v) donkey serum, and then incubated at 4°C overnight with the desired primary antibodies. They were washed 3× for 5 min with 1× PBS and incubated for 1 h with species-specific secondary antibodies (1:500) at room temperature. 6-Diamidino-2-phenylindole (DAPI, 1:1000) in deionized water was used to visualize the nuclei following secondary antibody. All images were captured with the TCS SP2 spectral confocal microscope (Leica Microsystems). Digital images were cropped and arranged using Photoshop CS (Adobe Systems).

### Assessment of the uptake of fluorescent beads by taste cells

Mice anesthetized by an intraperitoneal injection (10ml/kg) of a mixture of ketamine (4.28 mg/ml), xylazine (0.86 mg/ml), and acepromazine (0.14 mg/ml), were administered fluorescent polystyrene latex nanoparticles (Fluoresbrite YG; 200 nm diameter) (Polysciences, Warrington, PA; cat. No. 09834-10). After 2 hours, tongues were isolated and fixed using 4% paraformaldehyde in 1× PBS. Then, frozen sections of 10 μm thickness were obtained as described above and stained with an antibody specific for T1R3. Confocal images were captured as described above. Quantitative analysis of fluorescent beads uptake from lingual epithelium containing CVP was done as previously described(31). Analysis of the degree of bead uptake from oral cavity containing CVP by threshold analysis using ImageJ. Images of the fluorescent beads were saved as 8-bit grayscale images and then converted to binary images. The percentage of the pixels with a signal intensity that exceeded the threshold cutoff point of 75 out of 255 was calculated for the area occupied by CVP. Images of DAPI-staining nuclei in the same field acquired in a separate channel were threshold at a cutoff point of 70. The extend of bead uptake was expressed as the ratio of pixels with fluorescent beads to pixels with DAPI after normalization to a mean value of 1.0 for CVP from WT mice.

### Brief-access taste tests

Brief access tests were conducted using the Davis MS-160 mouse gustometer (Dilog Instruments, Tallahassee, FL) as described (73, 75, 76). The following taste compounds were tested: sucrose (30, 100, 300, 1000 mM), sucralose (1, 3, 10, 30 mM), monosodium glutamate (MSG; 30, 100, 300, 100 mM), denatonium (0.3, 1, 3, 10 mM), citric acid (1, 3, 10, 30 mM), and NaCl (30, 100, 300, 600 mM). Mice were water- and food-restricted (1 g food and 1.5 mL water) for 23.5 h before test sessions for appetitive taste compounds (sucrose, sucralose, and MSG). For the aversive taste compounds (citric acid, denatonium, and NaCl), mice were water- and food-deprived for 22.5 h before testing. In each test session, four different concentrations of each taste compound and water control were presented in a random order for 5 s after first lick, and the shutter reopened after a 7.5-s interval. The total test session time was 20 min. An additional 1-s “washout period” with water was interposed between each trial in sessions testing aversive tastants. *Spib^KO^* and their littermates *Spib^+/+^* mice were tested at the same time in parallel. Each mouse was tested with all the compounds. After each session mice were allowed to recover for 48 h with free access to food and water. Body weight of the mice was monitored daily, and the mice at or over 85% their initial were used. The ratio of taste stimulus to water licks was calculated by dividing the number of licks for taste compounds by the number of licks for water presented in the parallel test session. Lick ratios > 1 indicate preference behavior to the taste compound, and lick ratios < 1 indicate avoidance behavior to the taste compound.

### Statistical analyses

Prism (GraphPad Software) was used for statistical analyses, including calculation of mean values, standard errors, and unpaired t-tests of cell counts and qPCR data. Data from taste behavioral tests were compiled using Microsoft Excel. For statistical analyses of behavioral, two-way ANOVA and post hoc t-tests were used to evaluate the difference between genotype and concentration using OriginPro (OriginLab). *p*-Values < 0.05 were considered significant.

## Acknowledgements

This study was supported by NIH-NIDCD grants R01 DC014105 and P30 DC011735 to R.F.M. and Young Scientists Fund of the National Natural Science Foundation of China (31800875) to Y.Q. Imaging was performed at the Monell Histology and Cellular Localization Core, which is supported, in part, by funding from NIH-NIDCD Core Grant P30DC011735 and National Science Foundation Grant DBI-0216310 (to Gary Beauchamp). The authors wish to thank Dr. Lee Garrett-Sinha for providing the *Spib* knockout mouse strain and the SPIB antibody, Dr. Hong Wang for creative suggestions and Mr. Kevin Redding for technical assistance.

